# Effects of Experimental Autoimmune Encephalomyelitis on Bladder Function and Properties of Pelvic Ganglion Neurons

**DOI:** 10.64898/2025.12.04.691436

**Authors:** Sherryl L. Henderson, Virginia Garcia, David J. Schulz

**Author notes:** Contact: David J. Schulz.

## Abstract

Multiple sclerosis (MS) has substantial impacts on autonomic function. In part, MS results in loss of normal autonomic activity that contributes to disease-associated pathology such as neurogenic bladder, bowel, and sexual dysfunction. Yet little is known of the impacts of MS on peripheral autonomic neurons that directly innervate these target organs. In this study, we measured changes in properties of neurons of the mouse major pelvic ganglion (MPG) associated with two different levels of severity in mice with experimental autoimmune encephalomyelitis (EAE), a model of MS. Our data show that bladder function and physiological properties of MPG neurons are altered in association with EAE, and differ between less and more severe clinical scores associated with disease progression. EAE results in decreased bladder function measured by decreased urine output and increased micturition pressures. In the most severely affected animals, action potentials (APs) in MPG neurons show increased half-widths as a result of decreased maximum decay slopes leading to overall broader and longer APs and lower maximum firing frequency. These changes in AP properties are associated with differences in ion channel subunit expression as well. *KCNA1-4, KCNN3* and *SCN2A1 and SCN3A* mRNA levels in MPGs decreased with disease severity. Taken together, our data indicate that peripheral autonomic neurons are fundamentally altered in EAE, suggesting that longer-term therapeutic approaches for MS could target these neurons directly to potentially help ameliorate neurogenic target organ dysfunction.

## 1 INTRODUCTION

Multiple sclerosis (MS) is a chronic autoimmune disease characterized by inflammation, demyelination, and neurodegeneration in the central nervous system (CNS) (1; 2). A common and debilitating symptom of MS is neurogenic lower urinary tract dysfunction (LUTD), which affects over 90% of patients as the disease progresses (3; 4; 5). LUTD in MS manifests as detrusor overactivity, urinary retention, and detrusor-sphincter dyssynergia (DSD), significantly impairing quality of life and increasing the risk of urinary tract infections (6; 7; 8). While the central mechanisms of LUTD in MS have been studied, the role of peripheral autonomic ganglia, particularly the major pelvic ganglion (MPG), remains poorly understood.

The MPG is a critical autonomic hub that regulates lower urinary tract (LUT) function by integrating sympathetic and parasympathetic inputs to control bladder storage and voiding (9; 10; 11). In rodent models, MPG neurons exhibit plasticity in response to pathological conditions such as spinal cord injury (SCI) and diabetes, altering their excitability and synaptic properties (12; 13; 14). Experimental autoimmune encephalomyelitis (EAE), a widely used mouse model of MS (16; 17), replicates key features of MS-related LUTD, including bladder dysfunction (18; 19). However, the effects of EAE on MPG neuron properties and their contribution to LUTD remain unexplored.

This study investigates the impact of EAE on MPG neuron excitability, gene expression, and bladder function in female SJL/J mice. We hypothesize that EAE-induced CNS damage alters the input to MPG neurons, leading to changes in their intrinsic properties and contributing to LUT dysfunction. By combining urodynamic assessments, electrophysiology, and molecular analyses, we aim to elucidate the peripheral autonomic mechanisms underlying MS-related bladder dysfunction, addressing a critical gap in the current understanding of LUTD pathophysiology (20; 21).

## 2 METHODS

### Animals

All experimental procedures were approved by the University of Missouri Institutional Animal Care and Use Committee and followed the National Institutes of Health Guide for the Care and Use of Laboratory animals. The animals were kept in groups on a 12:12 hour light/dark cycle with food and water ad libitum. Adult female SJL/J were obtained from the Jackson Laboratory (Bar Harbor, ME). These adult female SJL/J were placed into three groups: control, EAE score 2, and EAE score 3.5.

### Induction of experimental autoimmune encephalomyelitis by active immunization

EAE was induced in female SJL/J 9 to 10 weeks of age by subcutaneous injection of 200uL proteolipid protein (PLP139-151) in complete Freund’s adjuvant (CFA) and ip injection of pertussis toxin (Ptx) obtained from Hooke Laboratories (St. Lawrence, MA). Mice underwent clinical observation and were scored daily to assess the progression of EAE. Assessments were reported using a standardized rating scale for the evaluation of motor deficits (17): 0 no deficit; 0.5 tip of tail is limp; 1 limp tail; 1.5 limp tail and hind leg inhibition; 2 limp tail and weakness of hind legs; 2.5 limp tail and dragging of hind legs; 3 limp tail and complete paralysis of hind legs; 3.5 limp tail and legs are together on one side of body; 4 limp tail, complete hind leg and partial front leg paralysis; 4.5 complete hind leg, and partial front leg paralysis with no movement around cage; 5 moribund. EAE in SJL mice follows a relapsing remitting time course of paralysis. In EAE induction by active immunization with PLP139-151 mice typically reach a score of 3 to 4.5 10-16 days post-induction. In this study mice were monitored for signs of urinary dysfunction and motor deficits starting 8 days post-induction and harvested at scores of 2 or more (Figure 1). A total of 45 control and EAE animals were used.

**Fig. 1.**
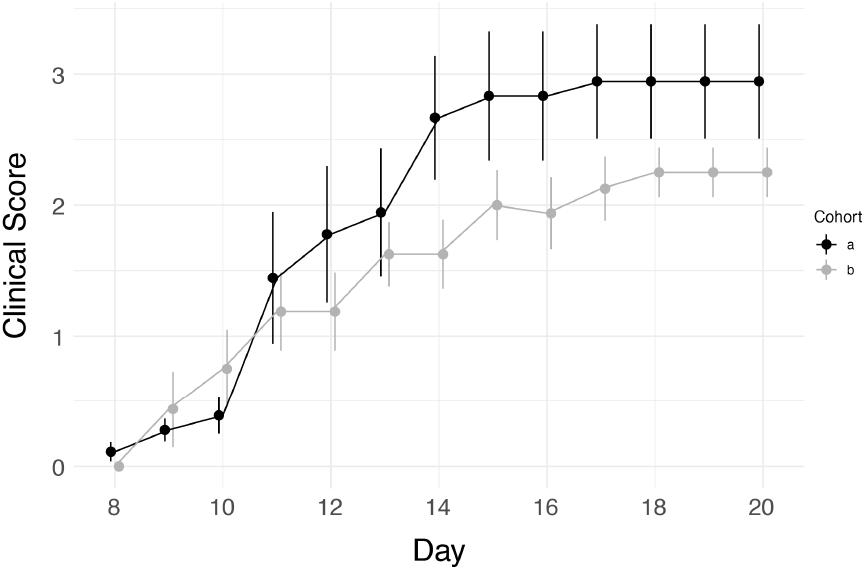
Time course of onset of symptoms associated with EAE in two cohorts of mice. Mice were induced in two experimental cohorts over a 3 to 5-day period. Both cohorts exhibited similar progression in clinical scores; however, animals in Cohort A reached a higher peak severity relative to Cohort B. For data analysis, animals were grouped based on their clinical scores across cohorts to facilitate comparison across disease severity levels.

### Tissue isolation for electrophysiology recordings

Mice were euthanized with overdose of isoflourane by inhalation and their MPGs were dissected out. The MPGs were pinned down in a Sylgard dish containing oxygenated physiological saline solution composed of, (in mM) NaCl, 146; KCl, 4.7; MgSO4, 0.6; NaHCO3, 1.6; NaH2PO4, 0.13; CaCL2, 2.5; Glucose, 7.8; HEPES, 20, adjusted to a pH of 7.3 (Jobling and Lim 2008). The saline was continuously perfused into the dish for in-vitro intracellular recordings.

### Electrophysiology recordings

Dissections and experiments were performed in oxygenated physiological saline at room temperature. The MPGs from 10–12-week-old female SJL/J were dissected and pinned in a Sylgard dish. The preparations were then de-sheathed by removing fat and connective tissue. Sharp electrodes pulled on a P-97 microelectrode puller (Sutter, Novato, CA) were filled with 500 mM KCl, and had resistances of 130 to 200 MOhm. Recordings were obtained using an Axoclamp 900A (Molecular Devices, Sunnyvale, CA) amplifier, and digitized at 10 kHz using a Digidata 1440 (Molecular Devices). Action potential properties, passive properties, and firing rate were calculated using the auto-statistics package of the Clampfit program from the pClamp 10.3 suite of software (Molecular Devices) and Spike2 (Cambridge Electronic Design Limited, Cambridge, England).

### Passive, excitability, and action potential properties

Resting membrane potential, input resistance, membrane time constant, membrane capacitance, rheobase, negative rheobase, max firing rate, halfwidth, antipeak, peak, max rise slope, max decay slope, rise slope, and decay slope were estimated from current injection step protocols from-300pA to 400pA by steps of 50pA or-500pA to 700pA by steps of 100pA current injections having a duration of 1000 ms.

### Void Spot Assays (VSAs)

Testing was performed in the vivarium where mice were housed. Individual mice were placed in chromatography paper-lined enclosures daily for 2 hours between 2 pm and 4 pm CST. Enclosures consisted of inverted polycarbonate mouse cages placed on a rack in the same animal facility room where the mice were housed. The mice could walk directly on the filter paper within the enclosure. The mice did not have access to food and water during testing. Seven-week-old mice were acclimated to testing by performing the VSAs for ten days before induction with EAE. Testing then was resumed four days post-induction and continued until peak illness (approximately 10-15 days). The filter paper was imaged under ultraviolet light using an iPhone 13 camera (Apple, Cupertino, CA) and UV flashlight source (uvBeast, Portland, OR). Filter paper that had been chewed thoroughly or had visibly overlapping spots was excluded. A standard curve was uploaded to Void Whizzard (29) so that the pictures could be analyzed. The data we obtained from the VSAs included void area, number of voids, and void volume. Void volume was determined by area using a standard curve generated by applying known volumes of liquid to the filter paper and imaging as above.

#### Cystometry

Before cystometric study, all animals were kept in standard cages with free access to food and water and normal 24hr light cycling. Mice were anesthetized by intraperitoneal injection of 1.2g/kg urethane. Supplemental doses were given to maintain an adequate plane of anesthesia. Mice were placed on a heating pad to maintain body temperature. Surgical scissors made a midline incision of about 1 cm in the lower abdominal area. The bladder was exposed carefully without damage to the surrounding tissues. A hole was placed in the dome of the bladder using a 25 gauge needle. A catheter was placed within the dome of the bladder and held in place using a purse string suture. The catheter was connected to a T-shaped stopcock, an infusion pump NE-300 Infusion Syringe P (New Era Pump Systems, Farmingdale, NY), and a PX600I pressure transducer (Edwards Lifesciences, Irvine, CA). 0.9% NaCl solution was infused into the bladder at a rate of 20 µl/min for one hour as the mouse acclimated to the catheter. If no voiding contractions were observed during 20 minutes of habituation, the flow was increased by 10 µl/min every 5 minutes until voiding contractions were observed. Animals were allowed to complete 10 voiding cycles and the last 5 were taken for analysis. Data were acquired using an HP 78534C Monitor/Terminal (Palo Alto, CA) and digitized via molecular devices Digidata 1440A to be recorded in Clampex 10.7 (Molecular Devices). Parameters assessed included: i) Rate 1 (the initial rate of infusion to induce a voiding contraction), ii) baseline pressure (average pressure between voids), iii) maximum pressure (peak pressure during cystometrogram), iv) micturition pressure (pressure during voiding contraction minus baseline), v) void duration (time of one complete void cycle), and vii) intercontraction interval. Mice with areflexic bladder or where holes were discovered in their bladders upon infusion were excluded from this study.

### Real-time quantitative polymerase chain reactions (qPCR)

Following electrophysiology experiments, whole MPGs from both left and right side of each animal were collected in 750 µl TRIzol (Invitrogen, Waltham, MA) for RNA extraction. Total RNA was isolated from MPGs according to the manufacturer’s protocol. cDNA was generated from 100 ng total RNA primed with a mixture of oligo-dT and random hexamers that was reverse transcribed using qScript reverse transcriptase (QuantaBio, Beverly, MA). The final volume of the reverse transcription reaction was 20 µl and contained a final concentration of 2.5 ng/µl random hexamers, 2.5 µM oligo-dT, 40 U of RNase inhibitor, and 200 U of reverse transcriptase. Following heat inactivation of the enzyme samples were diluted in ultrapure water to a final volume of 175 µml before this template was used in qPCR analyses.

cDNA was used in qRT-PCR reactions to quantify relative expression of Na+ and K+ channel subunits from paired whole MPGs. Primer sets were previously validated in this system (23; 24; 13). qPCR reactions consisted of primer pairs at a final concentration of 2.5 µM, cDNA template, and SsoAdvanced SYBR mastermix (BioRad, Hercules, CA) according to the manufacturer’s instructions. Reactions were carried out on a CFXConnect machine (BioRad) with a three-step cycle of 95ºC-15s, 58ºC-20s, 72ºC-20s, followed by a melt curve ramp from 65ºC to 95ºC. Data were acquired during the 72ºC step and every 0.5ºC of the melt curve. All reactions were run in triplicates of 10 µl, and the average Cq (quantification cycle) was used for analysis. Reactions were normalized to a fixed amount of total cellular RNA (25; 26) and expression level for each individual sample was then calculated as a fold-expression level relative to the median Cq of the control group as follows: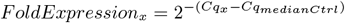.

### Statistics and Visualizations

Data were accumulated in Microsoft Excel then loaded into R version 4.5.0 for analysis (27). Visualizations were generated in R using the ggplot graphics package (28). ANCOVA assessed VSA trends over time. Wilcoxon Rank sum tests were performed to test statistical significance of cystometry parameters. Kruskal-Wallis tests followed by Dunn’s post hoc tests were performed to test statistical significance of passive properties, excitability properties, and action potential properties. Differences were deemed significant if *P* ≤.05.

## 3 RESULTS

### EAE Time course and Disease Progression

EAE was induced in twenty SJL/J mice in two batches (experiments a and b). EAE induction followed a relapsing-remitting disease course, with clinical symptoms appearing 8–10 days post-immunization and peaking at 10–16 days (Figure 1). Mice developed progressive motor deficits, including limp tails (score 1), hindlimb weakness (score 2), and complete hindlimb paralysis (score 3–3.5). Clinical scores plateaued by day 16–19, with no further progression before harvest. Two independent experimental cohorts exhibited similar disease trajectories, though the first cohort reached higher peak severity (Figure 1). Mice were grouped for analysis based on clinical scores (0 = control, 2 = moderate, 3.5 = severe), with bladder and MPG assessments conducted at peak illness. This time course aligns with established PLP-induced EAE models and confirms successful disease induction with associated neurological dysfunction.

### EAE alters urodynamics

VSAs were performed in all mice to determine if EAE altered the output of the lower urinary tract. End point cystometry was conducted with seventeen SJL/J mice to investigate urinary voiding dynamics. Daily urine spot volumes (Figure 2A) and counts (Figure 2B) were measured by void spot assay (VSA). Control animals showed consistent urine output throughout the testing period (Figure 2). In contrast, EAE animals exhibited variability in urine output, with a significant main effect of group on urine spot volume (ANCOVA, *F* (2, 233) = 5.260, *P* = 0.023; Figure 2). However, despite an overall downward trend across the time course of EAE progression no significant interaction effect between day and group was found. Cumulative void spot volumes (Figure 2A) and counts (Figure 2B) were significantly lower in EAE animals relative to control (*P <* 0.05).

**Fig. 2.**
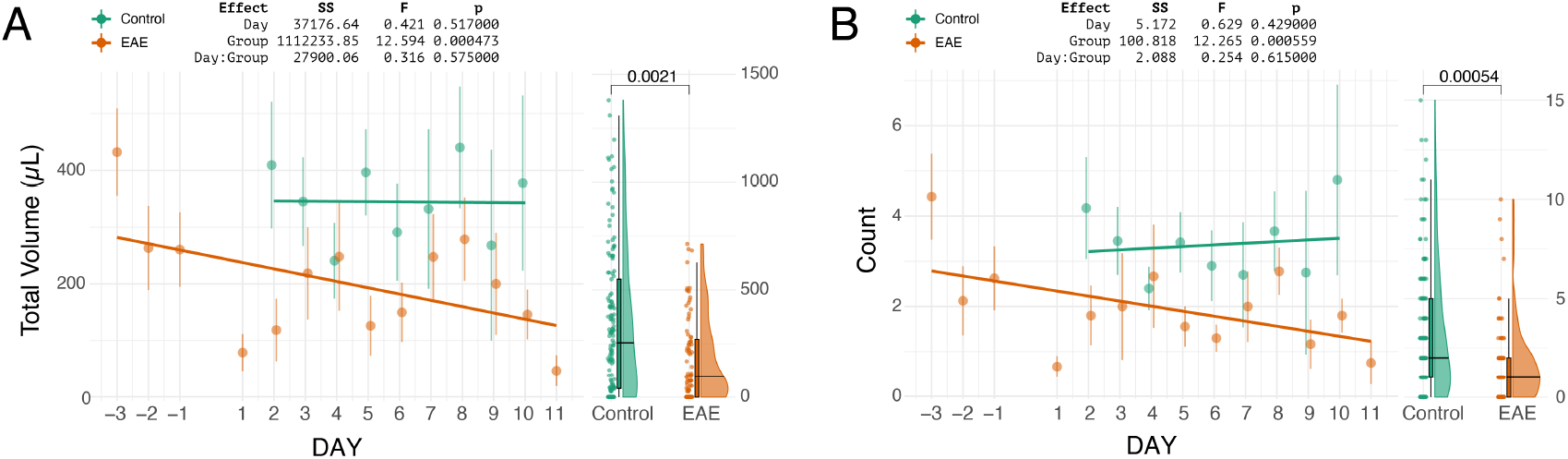
Daily urine output as measured by VSA in control and EAE animals. Measurements of (A) Volume and (B) Number of spots were made via VSA for a total of 40 animals. For EAE, the induction period and the subsequent post-induction period without testing are denoted by the gap in the x-axis. Days-3 to-1 represent the three days prior to induction and represent the control state for the EAE group, while days 1 to 11 correspond to the VSA data collected over 11 days following induction and EAE onset. Control animal data is displayed for a continuous 9 days, labeled as days 2 to 10. Tables represent result of ANCOVA. The cumulative volume and counts are also shown as mixed format plots that display raw data points, median and interquartile range (IQR) via boxplot, and distribution via half-violin plots. P-values on these plots refer to results of Wilcoxon tests.

**Fig. 3.**
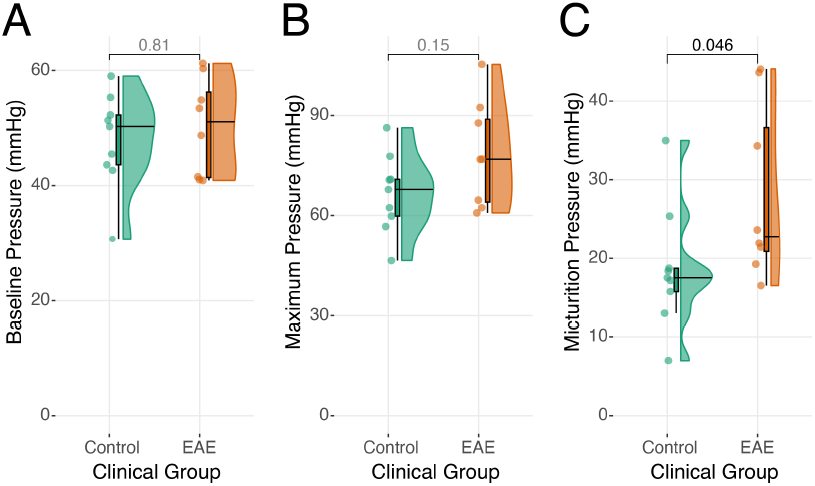
Effects of EAE on cystometry outputs. Plots showing cystometry data for baseline pressure (A), maximum pressure (B), and micturition pressure (C). The mixed format plots display raw data points, median and interquartile range (IQR) via boxplot, and distribution via half-violin plots. The was no significant effect of EAE on baseline pressure (*P* = 0.810) or maximum pressure (*P* = 0.149). However, micturition pressure was significantly higher in EAE animals relative to control (*P* = 0.049). The horizontal line within each violin plot also indicates the median. P-values shown as determined by Wilcoxon sign-rank tests.

Cystometry measurements revealed differences in bladder function between EAE and control groups, particularly in micturition pressure (3). Although no significant differences were observed in baseline bladder pressure (*P* = 0.810) or maximum pressure (*P* = 0.149) between the two groups, the micturition pressure during active voiding was significantly elevated in animals with EAE compared to controls (*P* = 0.049).

### Passive properties of MPG neurons remain unchanged in EAE animals

Intracellular recordings were performed in a total of 75 neurons across experimental animals grouped according to EAE clinical scores (0, 2, 3.5). Passive membrane characteristics resting membrane potential, input resistance, membrane time constant, and membrane capacitance were measured or calculated through systematic current injections employing two protocols:-300pA to +400pA steps) or-500pA to +700pA (100pA steps). No significant differences were detected in any of these passive properties in EAE animals at either level of clinical progression (Figure 4).

**Fig. 4.**
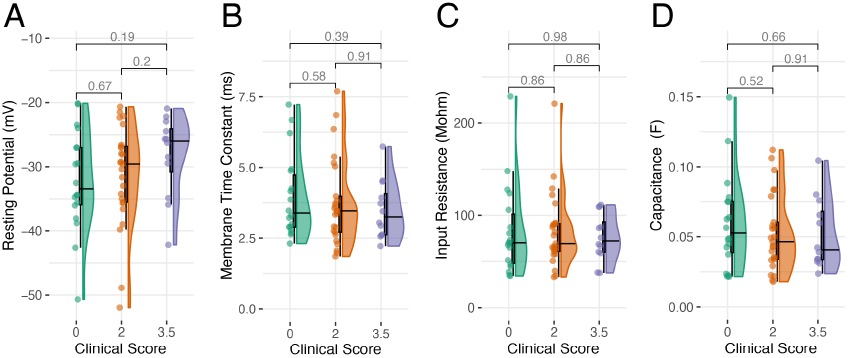
Effects of EAE on passive properties of MPG neurons. Plots showing intracellular recording data from neurons of the MPG for resting membrane potential (A), membrane time constant (B), input resistance (C), and capacitance (D). Plots as described in Figure 3. The were no significant effects of EAE on any of the passive properties measured in pelvic ganglion neurons. P-values shown for each comparison as determined by pairwise Dunn’s posthoc following Krusal-Wallis tests.

### EAE alters firing properties of MPG neurons

To assess the impact of EAE on firing properties of MPG cells, rheobase, negative rheobase, and maximum firing frequency were examined across clinical scores (Figure ref). While no significant differences in rheobase were observed between groups (*P >* 0.05; Figure 5), indicating that the minimum current required to elicit an action potential remained relatively stable regardless of disease severity, both negative rheobase and maximum firing frequency showed significant alterations in relation to clinical score. Negative rheobase, which reflects the minimum amount of current needed to elicit a rebound action potential when hyperpolarizing the neuron, was significantly reduced in more severe cases of EAE where clinical scores of 0 is significantly less than 3.5 (*P* = 0.002; 5). Additionally, maximum firing frequency, or the highest rate at which neurons can generate consecutive action potentials, was significantly altered by disease severity (Figure 5). In neurons from animals with a clinical score of 3.5, maximum firing frequency was reduced compared to score 0 (*P* = 0.013) and 2 (*P* = 0.026).

**Fig. 5.**
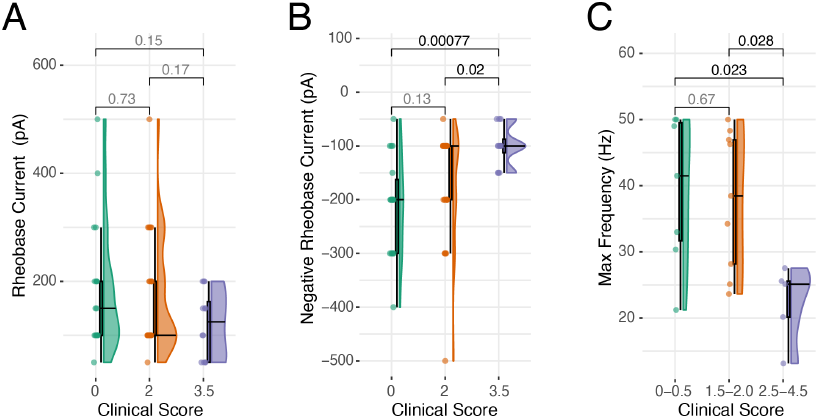
Effects of EAE on active properties of MPG neurons. Plots showing intracellular recording data from neurons of the MPG for rheobase (A), negative rheobase (B), and maximum firing frequency of tonic cells (C). Plots as described in Figure 3. The were no significant effects of EAE on rheobase (Kruskall Wallis test followed by Dunn’s posthoc). There was a significant decrease in negative rheobase in both EAE groups, as well as a decrease in the maximal firing frequency of the most advanced stages of EAE. P-values shown for each comparison as determined by pairwise Wilcoxon sign-rank tests.

### EAE alters action potential properties of MPG neurons

To assess the impact of EAE on MPG neuron excitability, action potential properties were characterized in seventy-five neurons. We characterized maximum rise slope, overall rise slope, maximum decay slope, overall decay slope, peak voltage, anti-peak (AHP) voltage, and half-width across clinical scores in the EAE model (Figure 6). The maximum rise slope and maximum decay slope were not significantly impacted by the severity of the disease (*P >* 0.05). Similarly, there were no significant differences in rise slope and decay slope, representing the rate of depolarization and repolarization (*P >* 0.05; 6). Additionally peak and anti-peak showed no significant differences across clinical scores (*P >*0.05). In contrast, half-width, a measure of action potential duration showed an overall significant difference (6). Post hoc analysis using Dunn’s test indicated that this significant difference was specifically driven by a comparison between the most severe clinical scores, with the half-width at a score of 3.5 being significantly greater than at a score of 2 (*P* = 0.043).

**Fig. 6.**
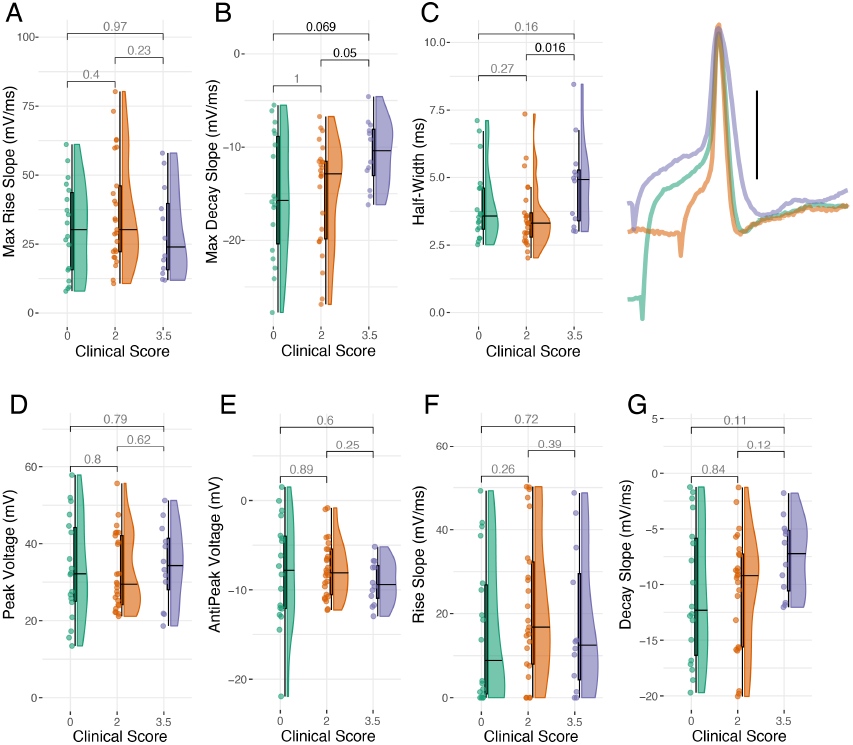
Effects of EAE on action potential properties of MPG neurons. Plots showing intracellular recording data of action potentials from neurons of the MPG. Plots as described in Figure 3. The were no significant effects of EAE on action potential peak voltage, anti-peak (AHP) voltage, or the rise slope of the action potential (Kruskall Wallis test followed by Dunn’s posthoc). There was a significant decrease in maximum decay slope, as well as an increase in the action potential half-width of the most advanced stages of EAE. P-values shown for each comparison as determined by pairwise Wilcoxon sign-rank tests. The inset shows representative action potentials overlaid from an individual neuron from each clinical score group. The duration of the traces is 10 ms and the vertical scale bar represents 10 mV.

### EAE alters expression of sodium and potassium channel subunits

We measured the steady-state abundance of mRNA in whole MPGs for three voltage-gated Na+ subunits (*SCN2A1, SCN3A* and *SCN7A*), the Kv1 and Kv2 K+ channel genes (*KCNA1-6* and *KCNB1-2* respectively), as well as calcium-activated K+ channel subunits for both SK-and BK-related currents (*KCNN1-3* and *KCNMA1*). Compared to controls, mice at the most severe levels of EAE progression (clinical scores 3.5) showed significantly decreased levels of *KCNA1-4* mRNA relative to both control and mice with a clinical score of 2.0 (*P >* 0.05; Figure 7). No differences in *KCNA5-6* or *KCNB1-2* were detected (Figure 7) across any groups. Further, the same differences were noted for only one of the SKKCa subunits, *KCNN3* (*P >* 0.05; Figure 8), while *KCNN1-2* and *KCNMA1* showed no differences (Figure 8). Conversely, all three Na+ channel subunits measured showed an increased level of expression in both EAE clinical score groups (2.0 and 3.5; *P >* 0.05; Figure 8).

**Fig. 7.**
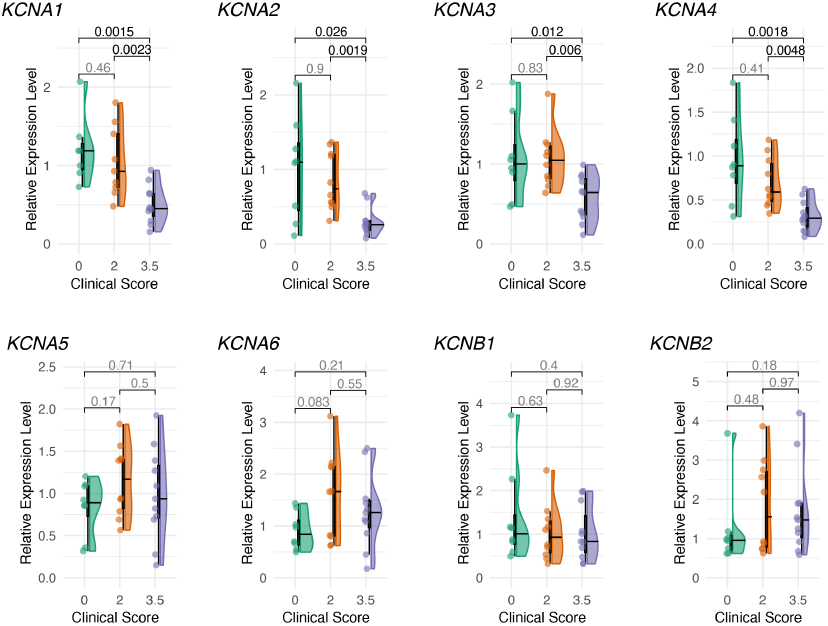
Effects of EAE on steady state mRNA levels of K+ channel subunits in the MPG. Relative expression levels of Kv1 (*KCNA*) and Kv2 (*KCNB*) potassium channel subunits were measured in whole paired MPGs from animals across three clinical scores (0, 2, 3.5). Each data point represents the paired MPGs from one animal. For each gene, each sample is expressed as a fold difference relative to the median Cq value of the control (clinical score 0) group. Plots as described in Figure 3. P-values are shown for each pairwise comparison are from Dunn’s posthoc tests following Kruskal-Wallis analysis.

**Fig. 8.**
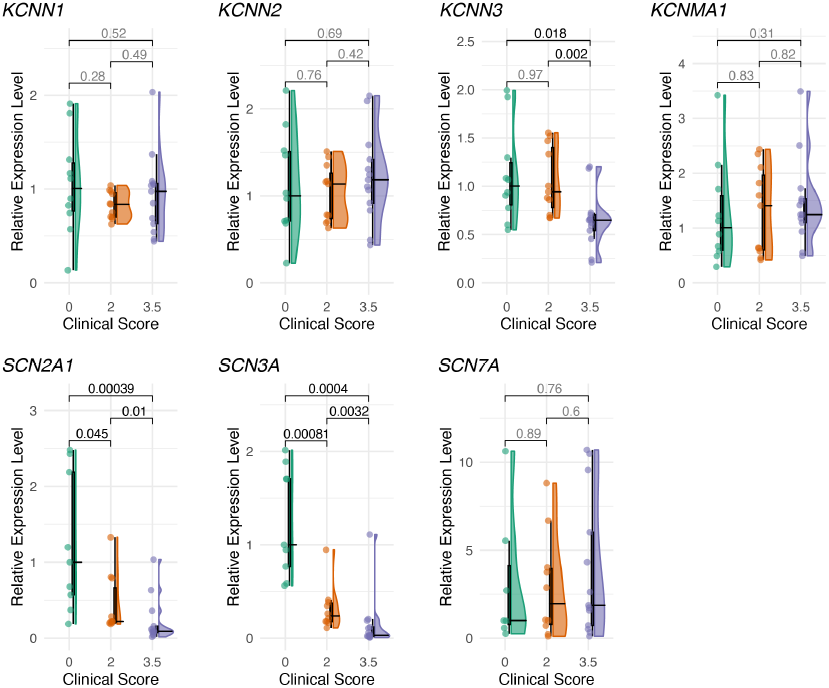
Effects of EAE on steady state mRNA levels of calcium activated-K+ channel and sodium channel subunits in the MPG. Relative expression levels of SKKCa (*KCNN*), BKKCa (*KCNMA1*) and Na+ (*SCN*) channel subunits were measured in whole paired MPGs from animals across three clinical scores (0, 2, 3.5). Each data point represents the paired MPGs from one animal. For each gene, each sample is expressed as a fold difference relative to the median Cq value of the control (clinical score 0) group. Plots as described in Figure 3. P-values are shown for each pairwise comparison are from Dunn’s posthoc tests following Kruskal-Wallis analysis.

## 4 DISCUSSION

Neurogenic bladder dysfunction is a prevalent and debilitating complication of multiple sclerosis (MS), affecting over 90% of patients as the disease progresses (3; 4). The disruption of neural pathways controlling micturition due to MS-related CNS lesions leads to phenotypes such as overactive bladder (OAB), underactive bladder (UAB), and detrusor-sphincter dyssynergia (DSD) (6). Our study, using the experimental autoimmune encephalomyelitis (EAE) model, replicates key features of MS-associated bladder dysfunction, including elevated micturition pressure and reduced urinary output, consistent with DSD (18; 19; 15). These findings align with clinical observations in MS patients, where DSD manifests as impaired coordination between bladder contraction and urethral sphincter relaxation, resulting in urinary retention and elevated post-void residuals (6; 21).

While central mechanisms of LUT dysfunction in MS have been more extensively studied, the role of peripheral autonomic ganglia, particularly the major pelvic ganglion (MPG), remains underexplored. The MPG integrates sympathetic and parasympathetic inputs to regulate bladder function (9), and its plasticity in response to pathological conditions like spinal cord injury (SCI) and diabetes is well-documented (13; 14). Our study reveals that EAE induces significant electrophysiological alterations in MPG neurons, including prolonged action potential duration (increased half-width) and reduced firing frequency, without affecting passive membrane properties. These changes suggest a selective disruption of active neuronal properties, likely mediated by ion channel dysregulation. This contrasts with diabetic models, where passive properties like resting membrane potential and input resistance are altered (14), suggesting distinct disease-specific mechanisms in EAE.

The downregulation of key ion channels subunits in EAE suggests a mechanistic basis for the observed electrophysiological changes. Potassium channel downregulation may result in delays in repolarization, manifest as a decrease in the maximum decay slope, resulting in prolonged action potential durations (30). The convergence of peripheral changes in MPG neuron excitability and central dysregulation in EAE mirrors the complex pathophysiology of MS-related LUT dysfunction. The elevated micturition pressure observed in our study could reflect compensatory detrusor contractions against a closed sphincter, a hallmark of DSD (31; 32; 33). Pharmacologically targeting the affected ion channels could offer therapeutic potential, as suggested by preclinical studies in other neurogenic bladder models (20).

Additionally, our findings underscore the need for early intervention to prevent irreversible autonomic dysfunction, given the progressive nature of MS (34; 35). A potential limitation of our study is the use of urethane anesthesia, which may transiently affect voiding reflexes (36). Future studies could employ awake cystometry to circumvent this issue. Moreover, investigating whether these neuronal changes persist during EAE remission or in progressive MS models would provide insights into the chronicity of autonomic dysfunction. Longitudinal assessments of MPG neuron properties and their correlation with bladder symptoms could further elucidate the temporal dynamics of LUT dysfunction in MS. Our study expands the understanding of MS-related LUT dysfunction by highlighting the critical role of peripheral autonomic plasticity. The hyperexcitable phenotype of MPG neurons in EAE, driven by ion channel dysregulation, complements central mechanisms and offers novel therapeutic targets. By bridging the gap between CNS pathology and peripheral autonomic dysfunction, this work paves the way for integrated approaches to manage neurogenic bladder in MS, ultimately improving patient quality of life.

## ACKNOWLEDGEMENTS

We would like to thank Dr. Steven Levine (KUMC) for invaluable guidance in learning the EAE induction protocols and related animal model advice. We also thank Madison Bunger for her help with organization of VSA data. This work was funded by a Pilot Grant from the National Multiple Sclerosis Society (PP-1804-30731).

